# The transcriptional regulator EarA and intergenic terminator sequences modulate archaellation in *Pyrococcus furiosus*

**DOI:** 10.1101/2023.06.15.545098

**Authors:** Richard Stöckl, Laura Nißl, Robert Reichelt, Reinhard Rachel, Dina Grohmann, Felix Grünberger

**Affiliations:** Institute of Microbiology and Archaea Centre, Faculty for Biology and Preclinical Medicine, University of Regensburg, Regensburg, Germany; Centre for Electron Microscopy, Faculty for Biology and Preclinical Medicine, University of Regensburg, Regensburg, Germany

**Keywords:** Archaea, Archaellum, transcriptomics, EarA, *Thermococcales*, single-molecule sequencing, transcriptional regulator

## Abstract

The regulation of archaellation, the formation of archaeal flagella-like appendages called archaella, is crucial for the motility, adhesion, and survival of archaeal organisms. Although the heavily archaellated and highly motile *Pyrococcus furiosus* is a key model organism for understanding the production and function of archaella in Euryarchaea, the transcriptional regulation of archaellum assembly is so far unknown. Here we show that the transcription factor EarA is the master regulator of the archaellum (*arl*) operon transcription, which is further modulated by intergenic transcription termination signals. *EarA* deletion or overexpression strains demonstrate that EarA is essential for archaellation in *P. furiosus* and governs the degree of archaellation. Providing a single-molecule update on the transcriptional landscape of the *arl* operon in *P. furiosus*, we identify sequence motifs for EarA binding upstream of the *arl* operon and intergenic terminator sequences as critical elements for fine-tuning the expression of the multicistronic *arl* cluster. Furthermore, transcriptome re-analysis across different *Thermococcales* species demonstrated a heterogeneous production of major archaellins, suggesting a more diverse composition of archaella than previously recognized. Overall, our study provides novel insights into the transcriptional regulation of archaellation and highlights the essential role of EarA in *Pyrococcus furiosus*. These findings advance our understanding of the mechanisms governing archaellation and have implications for the functional diversity of archaella.

## INTRODUCTION

Extracellular locomotive structures are ubiquitous across all domains of life and known as cilia or flagella in Eukaryotes, flagella in Bacteria, and archaella in Archaea (Khan and Scholey, 2018; Beeby et al., 2020). Though analogous in function to flagella, the archaellum is distinct in its evolutionary origin, genetic makeup, and the structural organization of its motility machinery (Jarrell et al., 2021; Nuno De Sousa Machado et al., 2022). Sharing similarities to the bacterial Type IV pili system, the archaellum distinguishes itself by its unique assembly and ATP-dependent rotation mechanism, in stark contrast to bacterial flagella, which operate based on ion gradients (Albers and Jarrell, 2015; Poweleit et al., 2016). Notably, in some archaea, the function of archaella extends beyond swimming and these cell appendages contribute to surface adhesion and biofilm formation through intra- and interspecies cell-cell contacts. Amongst other reasons, this functional diversity enables archaea to colonize a wide range of habitats, including extreme environments such as deep-sea black smokers (Näther et al., 2006; Schopf et al., 2008; Wirth, 2017; Wirth et al., 2018).

The complex architecture of the archaellum encompasses multiple components, including filament proteins (ArlA and/or ArlB), stator elements (ArlF, ArlG), proteins potentially needed for motor function (ArlC, ArlD, ArlE, ArlX), ATPase and ATPase-modulating proteins (ArlI, ArlJ), and platform proteins (ArlJ) (for an overview see (Jarrell et al., 2021)). Most of these components are encoded within the archaellum (*arl*) gene cluster. Arl gene clusters can be divided into the highly conserved Arl1 or Arl2 cluster that differ in their composition and phylogeny (Desmond et al., 2007). However, genes encoding associated proteins, such as enzymes for N-glycosylation of archaellins and chemotaxis system components, may be located in different regions of the genome (Schlesner et al., 2009; Jarrell et al., 2014).

To fine tune the expression of *arl* genes during different environmental conditions, diverse transcriptional regulation mechanisms have been discovered in Crenarchaeota (Jarrell et al., 2021). In *Sulfolobus*, transcription factor (TF)-based regulation has been extensively studied, revealing several factors that control archaellum formation. Transcription factors ArnA and ArnB play key roles in this process (Reimann et al., 2012). Other regulators that either impair (ArnC, ArnS, Abfr1, ArnR, ArnR1) or enhance (ArnD, PP2A) motility have also been identified (Lassak et al., 2013; Haurat et al., 2017; Hoffmann et al., 2017; Li et al., 2017). Notably, these proteins are absent in Euryarchaeota, where only a single regulator, the euryarchaeal archaellum regulator (EarA), was recently identified (Ding et al., 2016, 2017b).

This factor has been shown to be indispensable for transcription of the *arl* operon in *Methanococcus maripaludis*, which encodes most proteins involved in archaellum formation (Ding et al., 2016, 2017b). Interestingly, spontaneous mutation in the promoter region of an EarA knockout strain restored archaellum functionality (Ding et al., 2017a). Although the molecular mechanism was not revealed yet, it led to the hypothesis that EarA might aid in recruiting transcription factor B to a weak B recognition element (BRE) in the wild-type *arl* promoter, but may not be essential when the *arl* promoter has a strong BRE (Ding et al., 2017a). While a deletion mutant could be restored when EarA homologues from other archaea were heterologously expressed in *M. maripaludis*, this was not possible with the EarA from *Pyrococcus furiosus*, which could be attributed to lower sequence identity or codon usage issues (Ding et al., 2017b). This is particularly intriguing because among Euryarchaeota, *Pyrococcus* has served as a key model system to further our understanding of the archaellum machinery at the functional, structural, and regulatory levels (Näther et al., 2006; Näther-Schindler et al., 2014; Lewis et al., 2015; Daum et al., 2017; Chaudhury et al., 2018).

Given this discrepancy, our study investigated the role of the euryarchaeal archaellum regulator EarA in *P. furiosus*, finding that EarA is also essential for archaellation in the hyperthermophilic organism. Additionally, we present an updated single-molecule perspective of the transcriptional landscape of the *arl* operon. Finally, we contextualized our findings by integrating transcriptomics data from across the *Thermococcales* and examining the potential influence of sequence elements on the expression of the *arl* operon. This includes the presence of EarA binding sequences upstream of the *arl* promoter, as well as intergenic terminator sequences. Together these data suggest that while the transcription factor EarA is the key regulatory player of archaellum expression, intergenic sequences help in fine-tuning other components of the *arl* operon in *Thermococcales*.

## RESULTS

### The euryarchaeal archaellum regulator EarA is essential for archaellation in *Pyrococcus furiosus*

To elucidate the *in vivo* role of the regulator EarA, we took advantage of the existing genetic system of *P. furiosus* based on agmatine auxotrophy (parental wildtype) (Waege et al., 2010) and constructed a mutant strain (Δ*earA*) in which the *earA* gene locus including the corresponding promoter region was deleted. Gene rescue strains expressed EarA *in trans* either under the control of its native promoter (Δ*earA* recovery) or under the control of the strong, constitutive *gdh* promoter (Δ*earA* overexpression) (Micorescu et al., 2008). Successful deletion of the genomic *earA* locus was confirmed via PCR (Supplementary Figure 1A). Additionally, we assessed the recovery and overexpression of EarA levels within the Δ*earA* background by quantifying the production of archaellins using Western blot analysis (Supplementary Figure 1B,C).

The archaellation of these strains was visualized by transmission electron microscopy of Platinum/Carbon shadowed samples (Figure 1A-D, Supplementary Figure 2). The *earA* gene deletion led primarily to cells devoid of archaella, whereas overexpression resulted in heavily archaellated *P. furiosus* cells (Figure 1A-D). These outcomes mirrored the observed swimming behaviors, which indicated that no or few archaella in the deletion mutant are not sufficient to allow the cells to swim normally.

**Figure 1.**
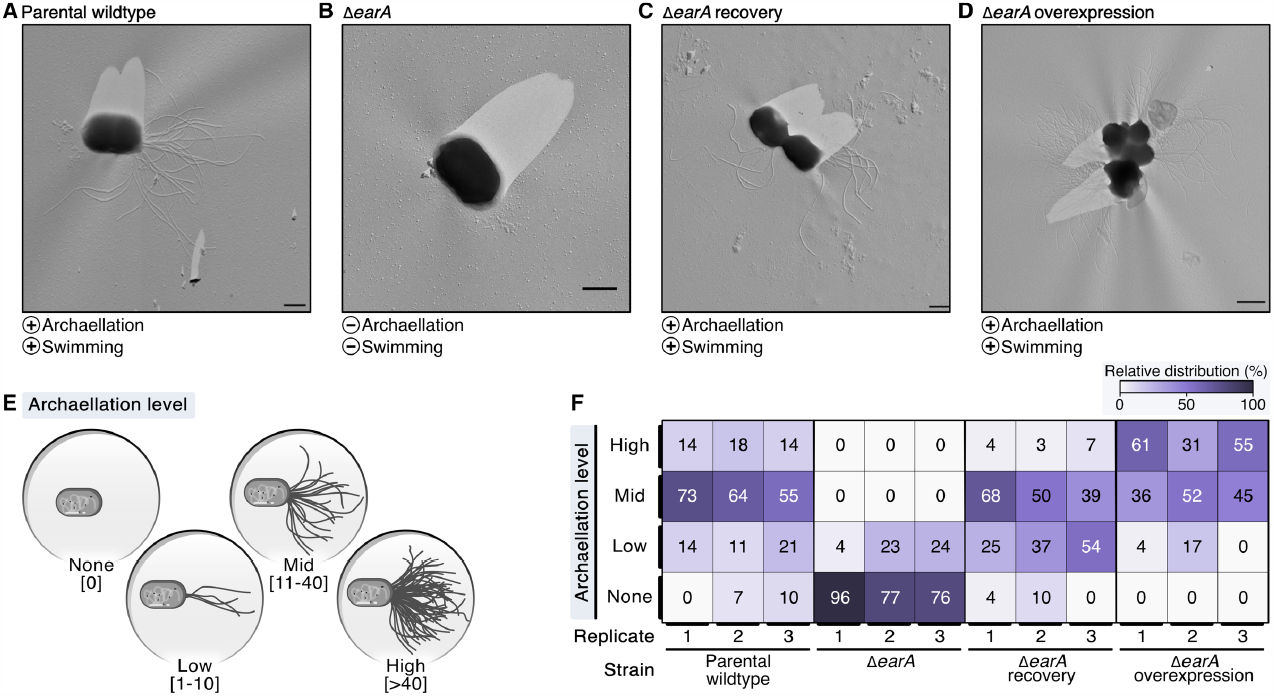
Electron microscopy evaluation of archaella formation in different genetic variants of *Pyrococcus furiosus* illustrates the central role of EarA. **A**, Transmission electron microscopy (TEM) micrographs of representative cells observed in the parental wildtype, **B**, Δ*earA* knockout strain, **C**, Δ*earA* recovery (native promoter) and **D**, Δ*earA* overexpression strain (*gdh* promoter). The presence and absence of archaellation and swimming behavior are denoted by plus and minus symbols, respectively. Scale bar: 0.5 μm. **E**, Systematic classification of the archaellation level based on the number of observable archaella originating from one cell. **F**, Color-coded representation (from 0% in white to 100% in violet) of the relative distribution of the four archaellation levels across the strains under investigation.

To systematically evaluate the observed initial differences in the extent of archaellation, we pursued further analysis by classifying the archaellation level based on the number of visible archaella of approximately 70-90 cells for each strain used in this study as none (0), low (1-10), mid (11-40) and high (>40) (Figure 1E and Supplementary Figure 2A). The increased EarA protein levels observed in the Δ*earA* overexpression strain indeed correlated with a significantly higher *in vivo* archaellation level clearly surpassing both the parental wildtype and the recovery strain (Figure 1F, Supplementary Figure 2B). These results underscore EarA as a pivotal regulator of archaellation in *P. furiosus*, suggesting that the cellular levels of this transcriptional regulator directly govern the degree of archaellation.

### A single-molecule update on the transcriptional regulation of the *arl* gene cluster in *P. furiosus*

To explore the potential hierarchic nature of EarA-induced transcriptional regulation, we analyzed the transcriptome landscape of the *arl* gene cluster in *Pyrococcus furiosus*. Therefore, we re-analyzed both short- and long-read RNA-seq data in addition to other database information, including transcription start and termination sites (Figure 2) (Grünberger et al., 2019, 2023). The Arl1-type cluster is characterized by three archaellin *(arlB0, arlB1, arlB2*) genes followed by the *arl* accessory genes *arlC, arlD, arlF, arlG, arlH, arlI, arlJ* and a hypothetical gene (Näther-Schindler et al., 2014; Jarrell et al., 2021). According to the DOOR2 database annotation, two genes preceding the *arl* genes, namely the transcriptional regulator EarA and the postulated methyltransferase *fam*, are part of the operon (Figure 2A,B). In contrast, prediction based on short-read RNA-seq data from mixed conditions suggests that the two genes are transcribed as an extra single unit (Figure 2A) (Mao et al., 2014; Grünberger et al., 2019).

**Figure 2.**
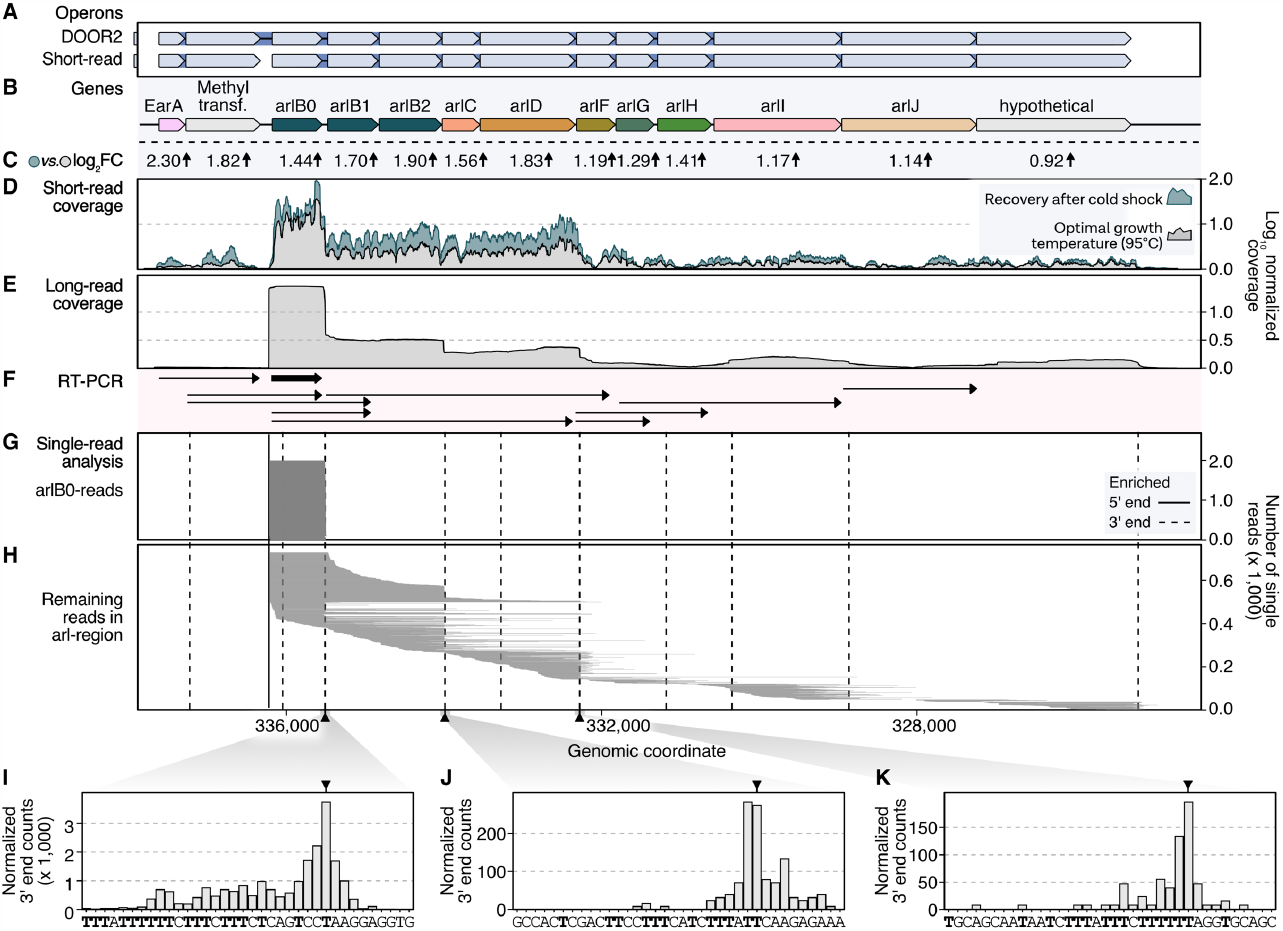
Long-read sequencing analysis provides high-resolution insights into transcriptional regulation of the *arl* gene cluster in *P. furiosus*. **A**, Annotated operons in the DOOR2 database and as suggested by ANNOgesic using short-read RNA sequencing of mixed growth conditions (Mao et al., 2014; Grünberger et al., 2019). **B**, Gene annotation of the *arl* cluster, including the transcriptional regulator EarA and a postulated methyltransferase upstream. **C**, Log2 fold changes obtained from differential gene expression analysis of samples recovered at 95°C after prolonged cold shock at 4°C (24 h) compared to cells in exponential growth phase (Grünberger et al., 2023). **D**, Mean coverage profiles normalized by CPM per position of these conditions are shown in grey (exponential phase) and blue (recovery) (Grünberger et al., 2023). **E**, Mean long-read sequencing coverage profile (Nanopore SQK-PCB109) of cells in exponential growth phase. **F**, RT-PCR results from (Näther-Schindler et al., 2014). Detected transcripts are visualized as arrows. *ArlB0* was detected as the major archaellin in *P. furiosus* and is highlighted in bold. **G**, Single-read plotting of reads mapping to *arl* region, split by reads mapping to *arlB0* and **H**, all other reads. Enriched primary 5’ ends from short-read sequencing of TEX-treated RNAs (Grünberger et al., 2019) and primary 3’ ends obtained from a recent Term-seq dataset (Grünberger et al., 2023) are shown as solid and dashed vertical lines, respectively. **I**, Histograms of CPM-normalized 3’ end positions in three regions (downstream *arlB0*: **I**, downstream *arlB2*: **J**, downstream *arlD*: **K**) that showed significant enrichment of 3’ ends using short-read Term-seq downstream of coding genes (Grünberger et al., 2023).

This was further supported by coverage re-analysis of short-read RNA-seq recently obtained from cells grown to exponential phase and cells recovered at the optimal temperature of 95°C following an extended cold shock at 4°C (Figure 2C,D) (Grünberger et al., 2023). Cells recovered from such conditions have been demonstrated to respond swiftly by swimming, which was confirmed by the induction of the *arl* gene cluster (Mora et al., 2014). Interestingly, we noted a considerable upregulation of the EarA regulator (log2 fold change: 2.3), initiating the activation of the *arl* genes. The level of activation decreased as the distance from the primary transcription start site increased.

To achieve a higher resolution of the transcriptional output, we re-analyzed long-read single-molecule data from samples cultivated under optimal growth conditions (Grünberger et al., 2023). Previous transcriptional start site mapping using differential RNA sequencing of Terminator-Exonuclease (TEX) treated RNA revealed that the primary start site is situated upstream of *arlB0*, the essential major archaellin protein (Grünberger et al., 2019). Indeed, the majority of the reads from long-read sequencing originate from this position. Furthermore, long-read coverage suggests differential expression of multiple transcription units, prompting us to refine and update the transcription model based on RT-PCR analysis (Figure 2E-H) (Näther-Schindler et al., 2014). In alignment with a previous study and structural data, our analysis verified that *arlB0* is the primary archaellin transcript, particularly during early exponential growth (Näther-Schindler et al., 2014; Daum et al., 2017). However, *arlB0* can be detected as co-transcript with downstream genes but also as a single transcript. Notably, we discovered several shortened units derived from the primary *arl* operon transcript (Figure 2H). These units exhibited a subset of reads originating from the upstream position of *arlB0*. Moreover, we detected transcripts with downstream read starts, suggesting the occurrence of processing events. Our findings revealed that transcripts spanning from *arlB0* to *arlJ* were not detected, in agreement with the previous study (Näther-Schindler et al., 2014). Importantly, the absence of enriched 5’ ends in our dataset, including the TEX-enriched deep-sequenced RNA-seq data from cells under various conditions, strongly suggests that all transcripts likely originate from *arlB0*. The hierarchical gene expression changes observed in the cold-shocked recovery cells further support this conclusion.

To validate these findings, we performed additional analysis on enriched 3’ ends and conducted a terminator sequence analysis (Figure 2I-K). This analysis confirmed the observed coverage profiles and highlighted the presence of poly(T)-sequences after *arlB0, arlB2*, and *arlD*. Given that poly(T) sequences have been shown to lead to intrinsic termination, as well as leading to the recruitment of the recently identified termination factors aCPSF1, these sequences likely serve as intergenic transcriptional terminators fine-tuning the expression originating from *arlB0* (Maier and Marchfelder, 2019; Sanders et al., 2020; Yue et al., 2020; Li et al., 2021; Grünberger et al., 2023).

### Investigation of transcriptome data and sequence features in *Thermococcales* reveal a heterogeneous production of major archaellins

We expanded our study by reanalyzing RNA-seq data from various organisms within the order *Thermococcales* including *Palaeococcus pacificus, Thermococcus onnurineus, Thermococcus kodakarensis*, and *Thermococcus litoralis* to understand the transcriptional characteristics of the *arl* gene cluster (Lee et al., 2015; Zeng et al., 2020; Liang et al., 2021; Villain et al., 2021; Grünberger et al., 2023). While the overall Arl1-like operon organization is well conserved among these organisms, they exhibited variations in the number of *arlB* genes, ranging from two to five gene variants (Figure 3A). Broadening the genomic analysis, in all but six of the 33 *Thermococcales* reference genomes currently available in the NCBI genomes page, arCOG Hidden Markov Model (HMM) matches for the genes encoding for EarA and Arl proteins were found (Supplementary Figure 3, Supplementary Table 2). The transcriptional regulator EarA is highly conserved in *Thermococcales* and mostly oriented in the same orientation as the arlB cluster, with some exceptions (Supplementary Figure 3,4). Note, that the NCBI annotations for the six reference genomes without HMM matches do not contain archaellin annotations and three of these organisms were described as “non-motile” or “not-flagellated” (Canganella et al., 1998; Grote et al., 1999; Hensley et al., 2014).

**Figure 3.**
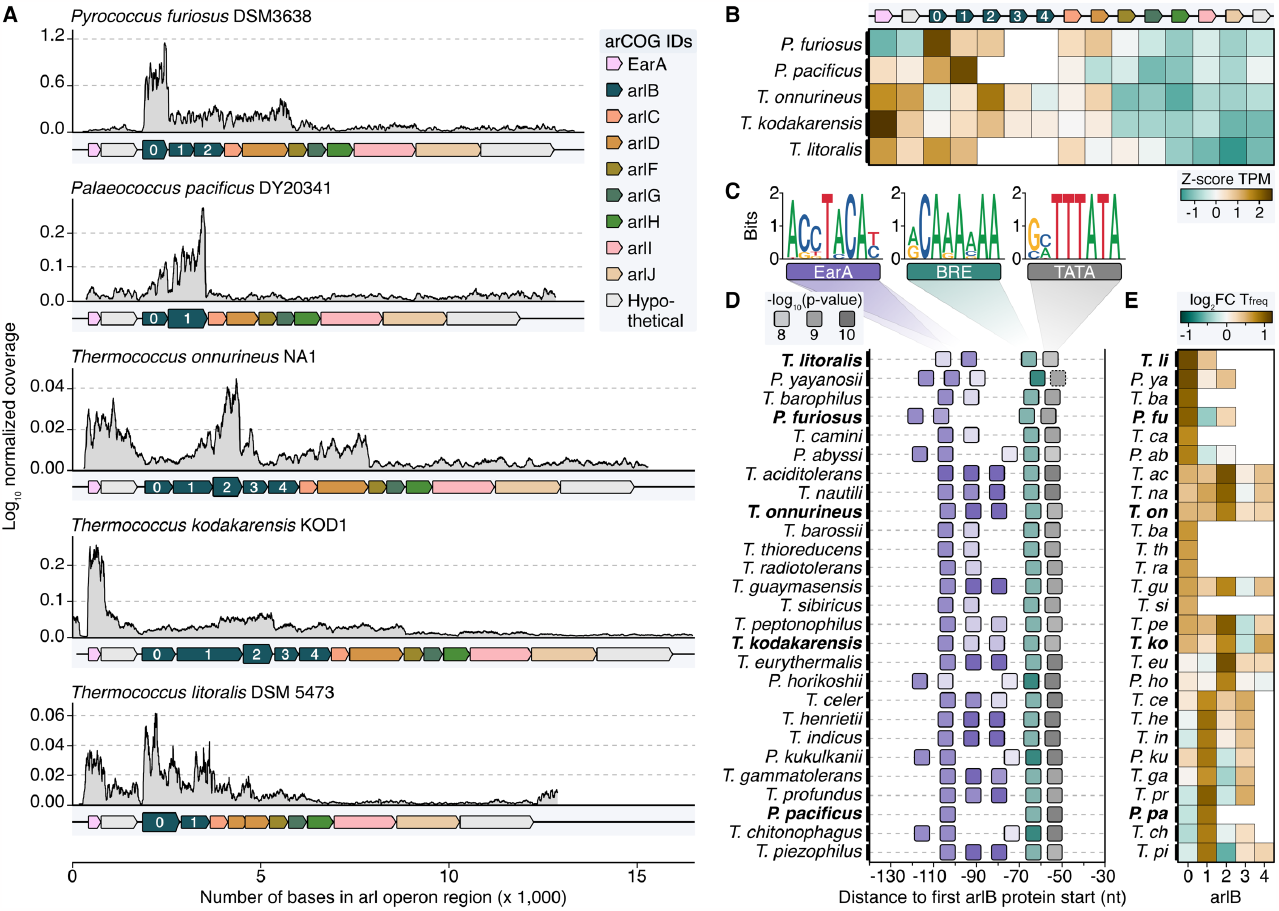
Transcriptome and sequence feature analysis reveal heterogeneous expression profiles of major archaellins in *Thermococcales*. **A**, Normalized coverage profiles from short-read sequencing of archaeal cells grown until exponential growth phase. Note that throughout the manuscript the most upstream *arl* gene was renamed *arlB0* in line with the current annotation in *P. furiosus*. Highest expressed *arlB* transcripts are highlighted in bold. **B**, Z-score normalized TPM values (brown: high TPM, green: low TPM) for all genes shown in A. **C**, Enriched motifs potentially recognized by EarA (pink), TFB (BRE, green) and TBP (TATA, grey) identified using MEME search (Bailey et al., 2009) of the sequence region 150 nt upstream of the first *arl* gene. **D**, Position of detected motifs (colored according to the description in C) are shown in a window from position -30 to -140 from the first *arlB* gene start. P-values are indicated by transparency of the boxes. Organisms shown in A and B are highlighted with bold text. The dashed box outline of the TATA box indicates that the element was not found by MEME but added after manual inspection. **E**, Enrichment of the base “T” 50 nt downstream of the respective *arlB* gene over genomic “T” frequency. Boxes are filled according to the log2 fold enrichment in brown (high “T”-frequency) or green (low “T”-frequency.

Comparing the coverage profiles of *Pyrococcus* with the other strains, we consistently observed a higher number of transcripts in the *arlB* region compared to the downstream accessory *arl* genes (Figure 3A). This finding indicates that under exponential growth conditions, *arlB* transcripts are the most abundant in all *Thermococcales*. In contrast to *Pyrococcus*, there were notable differences in the transcriptional output of the other strains. Notably, *arlB0* was not consistently the major archaellin based on expression levels. We observed some variability of the transcript levels, with either *arlB1* or *arlB2* as the most highly expressed gene or a balanced distribution between the *arlB* genes (Figure 2A,B). This balance was particularly evident for the five tandem co-transcribed archaellin genes in *T. kodakarensis*, aligning with previous Northern Blot results (Nagahisa et al., 1999). Also, the transcript counts of the regulator *earA* were significantly higher or equal to the *arlB* expression levels in the *Thermococcus* strains, suggesting it is transcribed as a separate unit.

Since previous analysis showed that EarA binds to a 6 bp consensus sequence (TACATA) in the promoter region of the first archaellin gene in *Methanococcus*, we performed sequence analysis in all *Thermococcales* (Ding et al., 2017b). We found that the *arl* loci in all organisms examined were preceded by archaea-typical BRE and TATA elements, which were uniformly located at a similar distance from the start codon of the first *arl* gene (Figure 3C,D). Specifically, the *arlB0* gene in *P. furiosus* featured a 30 nt long 5’ UTR (Grünberger et al., 2019).

Moreover, most cases showed a motif A(C/G)(C/G)TACA(T/C) at a fixed distance of nine nucleotides upstream of the BRE element, often repeating up to three times adjacently. We observed three distinct patterns, with either one (e.g., *P. horikoshii*), two (*P. furiosus*), or three EarA recognition sites (*T. kodakarensis*). It is noteworthy that the motifs in all *Pyrococcus* strains were shifted to either one window upstream or downstream compared to those in most of the *Thermococcus* strains. The absence of this motif in the upstream regions of all other *arl* genes supports the notion that EarA specifically binds upstream of the *arlB0* gene, thereby regulating the full gene cluster.

As a second layer of transcriptional regulation, we observed internal poly(T) sequences in *P. furiosus* (Figure 2I-K). Investigating whether this also applies to other *Thermococcales* and whether the strongest potential terminator sequence is positioned after the major archaellin, we calculated the enrichment of poly(T) sequences located 50 bases downstream of each *arlB* gene. Our results aligned with the overall expression levels, showing the highest T enrichment after the most abundant *arlB* gene (Figure 3E). The analysis across all *Thermococcales* confirmed our coverage analysis and indicated that, despite the *arl* gene organization, the most upstream *arl* gene is not always the most abundant gene, supporting the hypothesis that different ArlB might get incorporated into final archaellum filaments.

## DISCUSSION

In this study, we highlighted the crucial role of the euryarchaeal archaellum regulator EarA for archaellation within *P. furiosus*. Using TEM analysis of different genetic variants, we gathered *in vivo* evidence that the expression levels of EarA directly correlate with the level of archaellation. Given that the production of each protein component of the archaellum needs to be carefully controlled, we investigated how this control might be achieved on the transcriptional level (Chaban et al., 2007).

Our findings provide experimental support for prior speculations about EarA-based activation of transcription. Based on the occurrence of a spontaneous promoter mutation it was postulated that EarA might aid in the recruitment of TFB to a weak BRE element, marked by 7-8 consecutive adenines, in the *arl* gene cluster’s promoter (Ding et al., 2017a). This hypothesis is supported by our motif analysis which reveals a similarly weak BRE sequence present in all *Thermococcales* preceding the first *arl* gene. Notably, the approximated spacing of 9-11 base pairs between the first upstream located EarA binding site and the BRE implies a direct interaction of EarA with TFB. This interaction is reminiscent of the activation pattern observed for the potent transcriptional activator 2 (Ptr2), which is presumed to interact directly with TBP to recruit it to an imperfect TATA-box (Ouhammouch et al., 2005). Until now, only one archaeal transcriptional regulator, TFB recruitment factor 1 (TFB-RF1), that mediates TFB-recruitment, has been biochemically characterized in depth (Ochs et al., 2012). For both TFB-RF1 and EarA, it remains elusive which internal and/or external cellular stimuli impact their activities as transcriptional regulators (Reichelt et al., 2018). This aspect warrants more detailed examination, especially in organisms that encode both, the archaellum machinery and chemotaxis systems, such as *T. kodakarensis* (Briegel et al., 2017).

Moreover, transcription terminator-based regulation and the role of readthrough in archaella accessory gene production, have already been observed in *S. acidocaldarius* and appear to be important in *Thermococcales* as well (Lassak et al., 2012). Conceptually, this represents an elegant mechanism for the regulation of the multicistronic *arl* cluster allowing the fine-tuning of high-demand core components and lower-demand accessory genes (Thomas and Jarrell, 2001). However, extrapolating our findings to the final incorporation of the major archaellin into the archaellum is not straightforward, due to the potential influences of post-transcriptional and post-translational regulation. Intriguingly, our single-molecule analysis of transcripts in *P. furiosus* showed that mRNA stability could indeed play a role, potentially explaining why the most upstream gene is not always the highest expressed.

In the case of *Thermococcus kodakarensis*, RNA-seq data compare favorably with a quantitative proteomics study. The latter disclosed similar proportions (0.4 - 0.25 %) of total cell protein for *arlB0, arlB1*, and *arlB2*, hinting at a heterogeneous composition of the archaellum filament (Briegel et al., 2017). Although previous data were interpreted such that, like in Crenarchaeota, euryarchaeal archaella mainly consist of one type of archaellin only, emerging evidence contradicts this (Jarrell et al., 2021). Examples are the heterodimers of arlB1 and arlB2 observed in *Halorubrum lacusprofundi* (Pyatibratov et al., 2020) and *Methanocaldococcus villosus* (Gambelli et al., 2022). Whether this heterogeneity correlates with a structural organization of the archaellum analogous to the hook-, filament-, and cap-structures in the bacterial flagellum, or any type of multifunctionality of the archaellum, such as adhesion to cells or a variety of surfaces, warrants future investigation (Näther et al., 2006; Schopf et al., 2008; Weiner et al., 2012). Another intriguing prospect is exploring how the expression of a specific ArlB variant, presumably the principal archaellin, can be post-transcriptionally regulated if its encoding gene is not the most upstream *arlB* gene in the *arl* operon (*P. pacificus, T. onnurineus*).

In conclusion, our research uncovers new facets of the transcriptional regulation of the *arl* gene cluster in *P. furiosus* and establishes EarA as crucial transcriptional regulator for archaellation in *Pyrococcus furiosus*. Interestingly, the observed variation in *arlB* gene number and expression among *Thermococcales* indicates a more diverse production and assembly of archaellins than previously thought. Furthermore, our findings concerning the promoter motifs and intergenic poly(T)-sequences advance our understanding of the transcriptional regulation and potential mechanisms impacting the integration of *arl* genes into archaellum filaments.

## MATERIAL & METHODS

### Strains, Plasmids, Primers, and Antibodies

All strains, plasmids, primers, and antibodies used in this study are listed in Supplementary Table 1.

### Construction of vectors and transformation of *P. furiosus*

For the construction of *P. furiosus* strains Δ*earA*, Δ*earA* recovery and Δ*earA* overexpression, the modified genetic system for *P. furiosus* DSM3638 based on selection via agmatine auxotrophy as described in (Grünberger et al., 2021) was used.

For markerless disruption of the *P. furiosus earA* gene, the upstream and downstream flanking regions of the *P. furiosu*s *earA* gene including the promoter region were amplified using the primer pairs 0340upAscIFW/0340upFusRW and 0340doFW/ 0340doNotIRW (Supplementary Table 1). Both PCR products were used as a template for a single-overlap extension PCR and the resulting PCR product was cloned into the modified pMUR47 vector containing a two-gene resistance cassette via the AscI and NotI restriction sites (Grünberger et al., 2021). Successful and correct insertion of the PCR fragment into the vector was verified by DNA sequencing resulting in the plasmid pMUR443.

Circular plasmid DNA pMUR443 and strain MURPf37 were used for transformation and selection was carried out in 1/2 SME-starch liquid medium without agmatine sulfate and inosine+guanosine (I+G) at 85°C for 12 h (Grünberger et al., 2021). Pure cultures of the intermediate mutant MURPf76_i were obtained by plating the cells on solidified medium. The integration of the plasmid into the genome by single cross-over was verified by analyzing corresponding PCR products. For the counter selection cells were plated on solidified medium containing 50 μM 6-methylpurine and 8 mM agmatine sulfate to induce a second homologous recombination step to recycle the selection marker and to eliminate integrated plasmid sequences. The genotype of the final mutant (MURPf76) was confirmed by PCR and cells had to be grown in the presence of 8 mM agmatine sulfate and 8 mM I+G.

For EarA recovery and EarA overexpression in trans, a shuttle vector system for *P. furiosus* was used. Based on plasmid pMUR310 (Grünberger et al., 2021), an overexpression shuttle vector system was constructed that allows constitutive expression of a protein of interest via the *gdh* promoter (PCR amplified from *T. kodakarensis* genomic DNA using primers RPA_pYS13_GA_F/pYS13_GA-R) and the *hypA1* gene terminator (PCR amplified from *P. furiosus* genomic DNA using primers PYS14_GA_F/pYS13_RPA_GA_R) regions. Both regions were fused upstream and downstream of the gene of interest as described in (Waege et al., 2010) and cloned into pMUR310 via the EcoRV restriction site. One of these overexpression plasmids was used for PCR amplification of the plasmid backbone using primers pYS14Expgdh_F/pYS14_GA_F. The *earA* gene was PCR amplified from *P. furiosus* genomic DNA using primers flaaktivGAp914F/ flaaktivgap914r. Both PCR products were assembled using Gibson Assembly (Gibson et al., 2009) resulting in the plasmid pMUR444. Next, the Strep-tag® II sequence was inserted at the C-terminus via Phusion® High-Fidelity DNA Polymerase mutagenesis PCR (New England Biolabs) using the primers pYS14_GA_F/ 5’ phosphorylated flaakstrp14R to amplify the whole plasmid. The PCR product was ligated using T4 ligase (New England Biolabs) and transformed into *E. coli* DH5**α** cells (New England Biolabs), resulting in pMUR445. The final construct including the correct insertion of the Strep-tag® II sequence was verified by sequencing (Microsynth). Finally, to construct a shuttle vector expressing EarA under its native promoter the pMUR444 backbone including the *earA* gene, but without *gdh* promoter, was PCR amplified using primers flaaktivpromGAp914F/ GAp14bbminpromR. The *earA* promoter region was PCR amplified from *P. furiosus* genomic DNA using primers p14fusPF0340prom/ 034upRW1. Both PCR products were assembled using Gibson Assembly (Gibson et al., 2009) resulting in the plasmid pMUR458. The correct exchange of promoter regions was verified by sequencing (Microsynth).

1 μg of the circular plasmids pMUR310, pMUR445 and pMUR458 were transformed into MURPf76 as described previously (Waege et al., 2010; Kreuzer et al., 2013; Grünberger et al., 2021). Selection was carried out in 1/2 SME liquid medium without agmatine sulfate and I+G at 85°C for 12 h. Pure cultures of the mutants MURPf79 (Δ*earA* strain containing pMUR310), MURPf77 (Δ*earA* recovery strain containing pMUR458) and MURPf78 (Δ*earA* overexpression strain containing pMUR445) were obtained by plating the cells on solidified medium. Plasmid stability was verified by re-transformation into *E. coli* DH5**α** and DNA sequencing of purified plasmids (Microsynth). Final *P. furiosus* mutants could be grown in 1/2 SME medium (Fiala and Stetter, 1986) supplemented with 0.1% starch, 0.1% peptone and 0.1% yeast extract but without agmatine sulfate and I+G supplementation (Grünberger et al., 2021).

### Analysis of swimming behavior

*P. furiosus* strains were grown at 85°C in 1/2 SME medium (Fiala and Stetter, 1986) supplemented with 0.1% starch, 0.1% peptone and 0.1% yeast extract to a cell density of ∼1×106 cells per mL and swimming experiments were performed using a temperature gradient forming device (TGFD) at 100°C as described in (Mora et al., 2014).

### Western Blot analysis

*P. furiosu*s strains were grown at 85°C in 1/2 SME medium (Fiala and Stetter, 1986) supplemented with 0.1% starch, 0.1% peptone and 0.1% yeast extract to a cell density of ∼1×108 cells per mL and harvested by centrifugation at 20,500 g (4°C). The cell pellets from 20 mL cultures were resuspended in 1x PBS buffer and the cell suspensions were sonicated for 50 s using a Bandelin Electronic™ Sonopuls™ HD 3400 homogenizer (cycle: 80%, power 80%). After centrifugation for 30 min at 16,200 g (4°C), total protein concentrations of the cell extracts were quantified using the QubitTM Protein Assay Kit 3.0 Fluorometer (ThermoFisher). 2 μg of total protein extracts from each *P. furiosu*s strain were separated using SDS-PAGE. Western blots were performed as described previously (Waege et al., 2010).

### PCR test

*P. furiosu*s strains were grown at 85°C in 1/2 SME medium (Fiala and Stetter, 1986) supplemented with 0.1% starch, 0.1% peptone and 0.1% yeast extract to a cell density of ∼ 1×108 cells per mL and harvested by centrifugation at 20,500 g (4°C). Genomic DNAs were isolated using the ReliaPrep™ gDNA Tissue Miniprep System (Promega) following the Mouse tail protocol. The region of the *arlD* gene was PCR amplified using the primers Fladup_100fw/Fladdo_100rw, and the region of the *earA* gene locus was PCR amplified using the primers 0340do_113rw/0340up_100fw, respectively. PCR products were separated by agarose gel electrophoresis.

### Transmission Electron Microscopy

*P. furiosus* cells grown to the exponential phase were taken up in a syringe and carefully filtered through a polyamide/nylon filter (pore size 0.2 μm). The concentrated cell film was resuspended in 20-40 μL fresh media and chemically fixed with 1% (v/v) glutardialdehyde (final concentration) at room temperature. 5 μL of the fixed cell suspension were applied onto copper grids (400-mesh; G2400C, Plano) coated with a 10 nm carbon film (Carbon Coater 208 Turbo, Cressington), washed twice by blotting with distilled water, and the samples were subsequently shadowed with ∼1.0 nm Pt/C (15° angle; CFE-50, Cressington). Transmission electron micrographs were recorded on a CM12 transmission electron microscope (FEI) operated at 120 keV and fitted with a slow-scan CCD camera (TEM 0124; TVIPS) using EMMENU v4.0 (TVIPS).

### Gene calling and re-annotation

All currently available *Thermococcales* RefSeq reference genomes with a completeness of scaffold or better were acquired via the NCBI genome page (https://www.ncbi.nlm.nih.gov/data-hub/genome/?taxon=2258&reference_only=true&assembly_level=1%3A3; accessed 2023-05-12) (compare Supplementary Table 2).

The genomes were annotated via the arCOG definitions as described elsewhere (Dombrowski et al., 2020). In short, CDS sites and protein sequences predicted with Prokka (v. 1.14.6) (Seemann, 2014) were searched against arCOG (https://ftp.ncbi.nih.gov/pub/wolf/COGs/arCOG/zip.aliar14.tgz) (Makarova et al., 2015a) Hidden Markov Models (HMM) prepared with hmmbuild using hmmsearch (HMMER v. 3.3.2; Nov 2020; http://hmmer.org/). The best hit was assigned as annotation based on bit score then e-value, after an e-value cutoff <= 1e-3 (see Supplementary Table 2).

### Re-analysis of short-read RNA-seq data

For coverage analysis of the *arl* gene cluster the following RNA-seq dataset were reanalyzed: lllumina data from *Pyrococcus furiosus* DSM3638 exponential growth phase at the optimum growth temperature of 95°C (ERR11200497, ERR11200498, ERR11200499, ERR11200500) (Grünberger et al., 2023), *Palaeococcus pacificus* DY20341 cells from exponential growth phase with S0 addition (SRR10749090) (Zeng et al., 2020), *Thermococcus onnurineus* NA1 cells from exponential phase in CO-supplied media (SRR1702360, SRR1702361) (Lee et al., 2015), *Thermococcus litoralis* DSM5473 cells from exponential phase (SRR12486532, SRR12486546, SRR12486547) (Liang et al., 2021) and *Thermococcus kodakarensis* KOD1 cells from exponential phase (ERR6384078, ERR6384079, ERR6384080, ERR6384081) (Villain et al., 2021)

Sequencing reads in fastq format were filtered and trimmed using fastp (v. 0.23.3) to remove low-quality reads and adapter sequences (--cut_front, --cut_tail, --qualified_quality_phred 30, -- length_required 30) using parameters selective for single-end or paired-end reads (Chen et al., 2018). Subsequently, trimmed reads were aligned to the respective NCBI reference genomes (*P. furiosus*: NZ_CP023154.1, *P. pacificus*: NZ_CP006019.1, *T. onnurineus*: NC_011529.1, *T. litoralis*: NC_022084.1, *T. kodakarensis*: NC_006624.1) using Bowtie2 (v.2.5.1) with default parameters (Langmead and Salzberg, 2012). The resulting sequence alignment files (SAM) were converted to binary mapping format (BAM) and sorted using samtools (v.1.17) (Li et al., 2009). Additionally, position-specific coverage files were generated using samtools depth (-a -J), including reads with deletions in the coverage computation. Downstream normalization and visualization was performed using the Tidyverse in R (Wickham et al., 2019). Briefly, coverage for each position was first normalized by counts per million (CPM), before calculating the mean value of all replicates.

For calculation of transcripts per million (TPM) values of the protein-coding genes, featureCounts (RSubread package v. 2.10.5) was used to calculate the count matrix based on custom GFF files containing all *arl* genes detected using HMM as described above (Liao et al., 2019). Next, TPM values were calculated by dividing the number of reads mapping to each gene by the gene length in kilobases, then dividing the resulting reads per kilobase (RPK) values by the sum of all RPK values in the sample, and finally multiplying the quotient by one million. For visualization, the z-score was calculated for all genes in the *arl* gene cluster region, including the two upstream genes.

### Re-analysis of Nanopore RNA-seq data

Nanopore data from two biological replicates of *Pyrococcus furiosus* DSM3638 cells from exponential growth phase grown at the optimum growth temperature of 95°C were reanalyzed (ENA-project: PRJEB61177, runs: ERR11203080, ERR11203081) (Grünberger et al., 2023). Generation of the dataset, including detailed information about cell growth, RNA isolation, RNA treatment, library preparation, sequencing following the PCR-cDNA barcoding kit protocol (SQK-PCB109) from Oxford Nanopore Technologies (ONT) and data analysis is described in (Grünberger et al., 2023). Briefly, basecalling and demultiplexing was performed using guppy (v. 6.4.2+97a7f06) in high-accuracy mode. Full-length sequenced reads were detected, strand-oriented and trimmed using pychopper (v.2.7.2, https://github.com/epi2me-labs/pychopper) with autotuned cutoffs and the edlib backend for identifying the custom VN primer (5’-ACTTGCCTGTCGCTCTATCTTCATTGATGGTGCCTACAG-3’).

Mapping to the *P. furiosus* DSM 3638 genome (NCBI:NZ_CP023154.1) was performed using minimap2 (v. 2.24-r1122) with standard parameters suggested for aligning Nanopore genomic reads (-ax map-ont) (Li, 2018, 2021). Soft or hard clipped (>5 bases) reads were removed using samclip (v. 0.4.0), and SAM files were converted to sorted BAM files using samtools (v. 1.16.1) (Li et al., 2009). Coverage files generation including samtools depth and CPM normalization in R was performed as described for the short-read datasets. Single-read tracks of unspliced reads (njunc = 0) were plotted using ggplot2 geom_segment from read start to end.

### Sequence analysis and motif identification

Annotations were filtered using R and the Tidyverse packages (v. 2.0.0) (Wickham et al., 2019). Genomic sequences annotated as EarA were extracted and translated using seqkit (v. 2.4.0) (Shen et al., 2016). EarA protein sequences were aligned using ClustalO (v. 1.2.3) (Sievers and Higgins, 2014) with default parameters and visualized using ggmsa (v. 1.4.0) (Zhou et al., 2022).

The genomic locations that matched the “FlaB” HMM were annotated as “arlB” and assigned a rank, with the most-upstream location designated as “arlB0” to be consistent with the annotation used in *Pyrococcus furiosus*. Please note that this deviates from the database annotation of other strains that start with “arlB1”.

Genomic regions located 150 nt upstream of *arlB*0 and 50 nt downstream of all *arlB* genes were defined using plyranges (v. 1.18.0) (Lee et al., 2019). The sequences corresponding to these regions were extracted using seqkit subseq.

Motif analysis was conducted on the sequences upstream of the *arlB*0 genes using MEME (v. 5.5.2; -mod anr -dna -w 8) (Bailey et al., 2015). The identified motifs and their respective locations were visualized using the ggmotif (v. 0.2.1) (Li et al., 2022), ggseqlogo (v. 0.1) (Wagih, 2017), memes (v. 1.6.0), and Tidyverse packages.

The enrichment of “T” bases in the genomic regions downstream of the *arlB* genes was calculated as the log2 fold change of the letter frequency in the 50 nt downstream region, obtained using bedtools nuc (v. 2.31.0) (Quinlan and Hall, 2010), to the average letter frequency in the whole genome retrieved using the ‘letterFrequency(“T”)’ function from the Biostrings package (v. 2.66.0).

## AUTHOR CONTRIBUTIONS

RS performed bioinformatic analysis of the data, with contributions from FG. LS performed analysis of swimming behavior and TEM imaging. RRe performed construction of vectors. RRe, RRa and DG supervised this work. DG acquired funding to conduct the work. FG prepared the figures, with contribution of all authors. RS, RRe, DG and FG wrote the paper, with all authors contributing to manuscript revision, and approval of the submitted version.

## ACKNOWLEDGMENTS

We thank all the members of the Grohmann lab for fruitful discussions. Also, we would like to thank all current and previous members of the “archaeal transcription group” (especially Winfried Hauser, Zubeir El Ahmad and Martin Fenk) for establishment of the genetic system and sequencing protocols and Annett Bellack for providing antibodies.

## SUPPLEMENTARY FIGURES

**Supplementary Figure 1.**
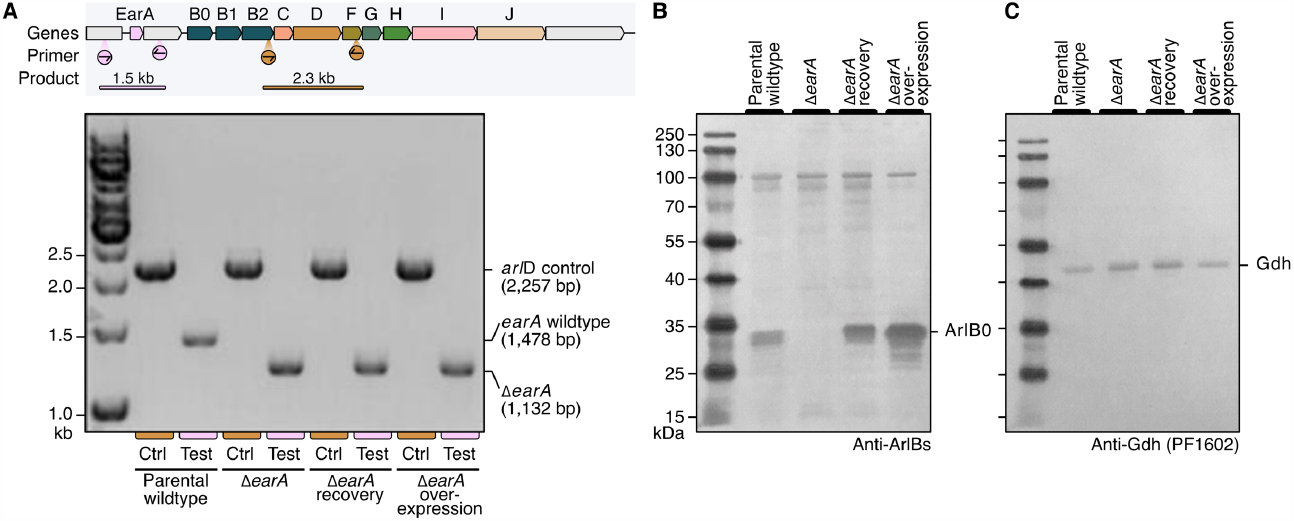
Confirmation of gene deletion and estimation of EarA expression levels in *P. furiosus* EarA deletion and recovery strains. **A**, Gene annotation of the *arl* gene cluster in *Pyrococcus furiosus*, with forward/reverse primers highlighted in circles (*arlD*: Fladup_100fw/Fladdo_100rw, *earA*: 0340do_113rw/0340up_100fw) and final PCR product in pink (*earA*) and orange (*arlD*). Results from PCR amplification of genomic DNA purified from the four strains used in this study using the two primer sets shown in panel A. **B**, Western Blot analysis of protein extract from the four strains used in this study using Anti-ArlB antibody and **C**, Anti-Gdh control.

**Supplementary Figure 2.**
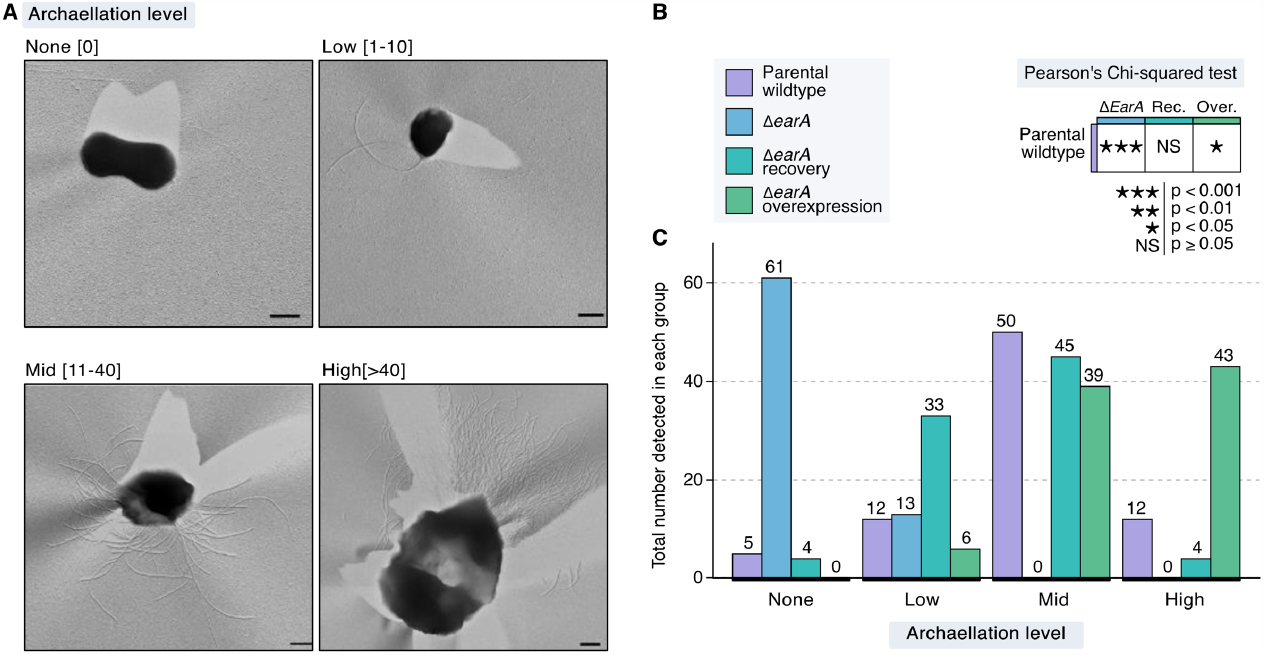
Systematic evaluation of archaellation in genetic *earA* variants. **A**, Representative TEM micrographs for the four levels of archaellation that were used to classify micrographs of the genetic variants, based on the number of visible archaealla as none (0), low (1-10), mid (11-40) and high (>40). **B**, Statistical test (Chi-squared test of mean number of observations in each level) to test significance of differential archaellation in the genetic variants. **C**, Total number of detected observations in each archaellation level color-coded by strain variant.

**Supplementary Figure 3.**
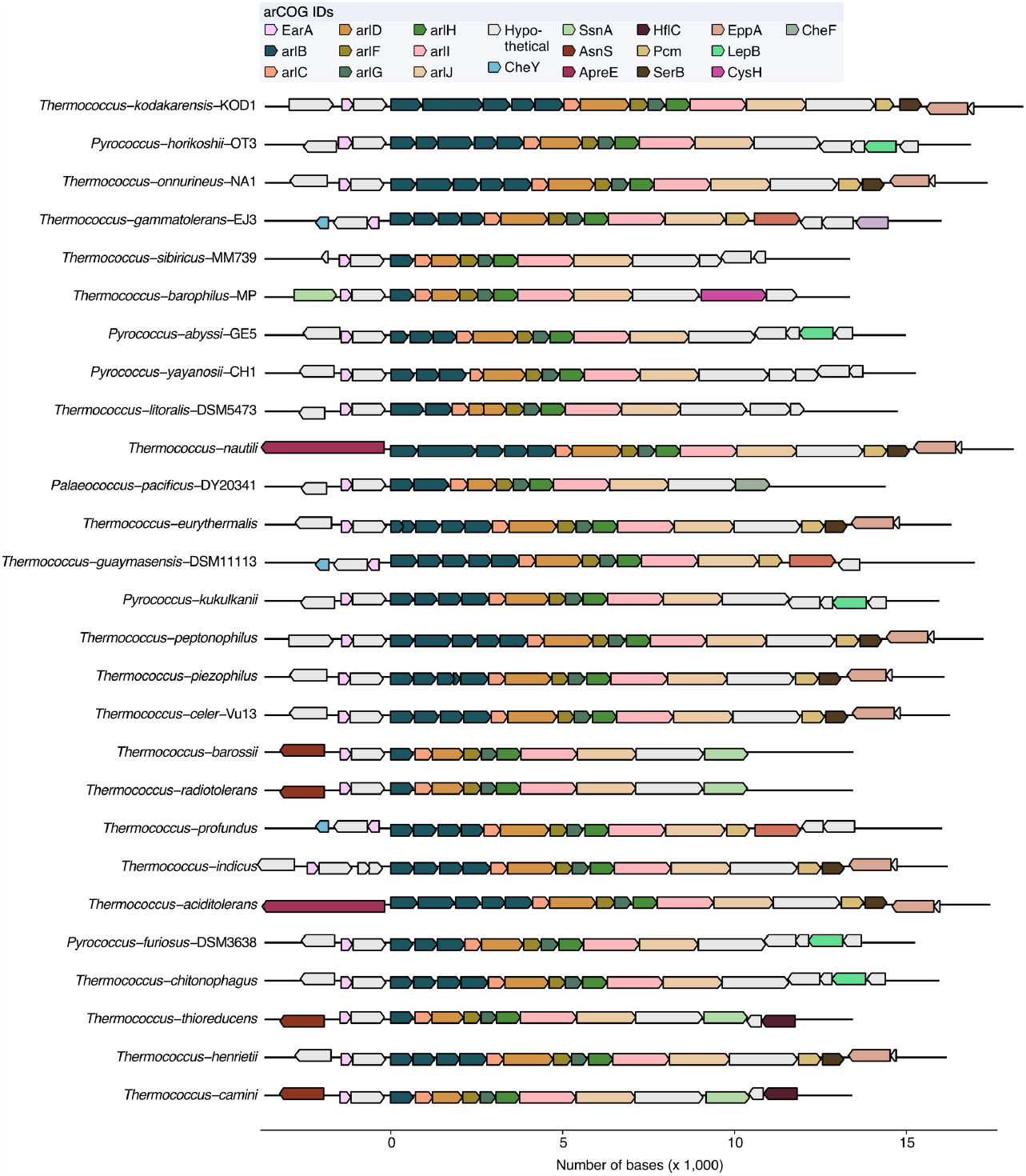
Genomic contexts of the *arl* gene clusters in *Thermococcales* species as defined by arCOG. HMM matches to protein sequences predicted by Prokka v. 1.14.6, centered on the *arlB0* gene (Seemann, 2014; Makarova et al., 2015b). Strandness is indicated by the arrow directions. All gene clusters exhibit a conserved structure comprised of single copies of the genes *arlC-J* preceeded by one to five *arlB* paralogs. The gene coding for the transcriptional regulator EarA is located immediately upstream of the *arl* gene cluster.

**Supplementary Figure 4.**
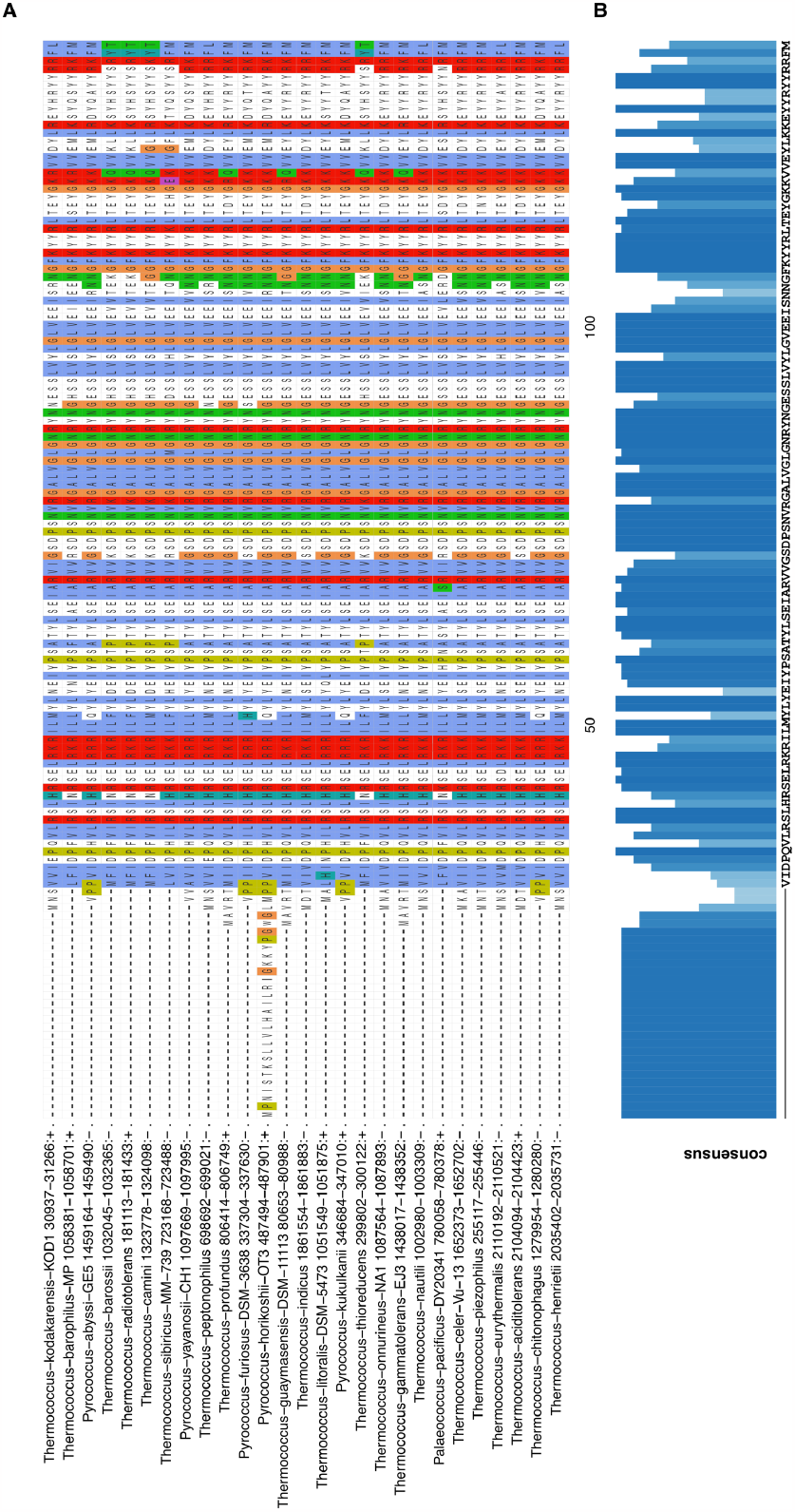
Multiple sequence alignment of EarA protein sequences in *Thermococcales*. **A**, Protein sequences predicted by Prokka v. 1.14.6 that matched to the EarA arCOG HMM were aligned using ClustalO (v. 1.2.3) (Seemann, 2014; Sievers and Higgins, 2014; Makarova et al., 2015b). Residues are colored based on their properties and identities: hydrophobic (blue), positive charge (red), negative charge (magenta), polar (green), cysteines (pink), glycines (orange), prolines (yellow), aromatic (cyan). **B**,Sequence conservation is indicated by the bar char and the resulting consensus sequence is given below. Genomic position and strandness of the respective gene is indicated next to the organism name. All EarA sequences exhibit highly conserved regions intersperced with mostly conserved regions.

